# Lateral compression of lipids drives transbilayer coupling of liquid-like protein condensates

**DOI:** 10.1101/2022.12.21.521462

**Authors:** Yohan Lee, Sujin Park, Feng Yuan, Carl C. Hayden, Siyoung Q. Choi, Jeanne C. Stachowiak

**Affiliations:** Department of Biomedical Engineering, The University of Texas at Austin, Austin, TX, USA; Department of Chemical and Biomolecular Engineering, Korea Advanced Institute of Science and Technology (KAIST), Daejeon, Republic of Korea; Institute for Cellular and Molecular Biology, The University of Texas at Austin, Austin, TX, USA

## Abstract

Liquid-liquid phase separation of proteins has recently been observed on the surfaces of biological membranes, where it plays a role in diverse cellular processes, from assembly of focal adhesions and the immunological synapse, to biogenesis of trafficking vesicles. Interestingly in each of these cases, proteins on both surfaces of the membrane are thought to participate, suggesting that protein phase separation could be coupled across the membrane. To explore this possibility, we used an array of freestanding planar lipid membranes to observe protein phase separation simultaneously on both surfaces of lipid bilayers. When proteins known to engage in phase separation bound to the surfaces of these membranes, two-dimensional, protein-rich phases rapidly emerged. These phases displayed the hallmarks of a liquid, coarsening over time by fusing and re-rounding. Interestingly, we observed that protein-rich domains on one side of the membrane colocalized with those on the other side, resulting in transbilayer coupling. How do liquid-like protein phases communicate across the lipid bilayer? Our results, based on lipid probe partitioning and the differential mobility of proteins and lipids, collectively suggest an entropic coupling mechanism, which relies on the ability of protein phase separation to locally reduce the entropy of the underlying lipid membrane, most likely by increasing lipid packing. Regions of reduced entropy then colocalize across the bilayer to minimize the overall free energy of the membrane. These findings suggest a previously unknown mechanism by which cellular signals originating from one side of the membrane, triggered by protein phase separation, can be transferred to the opposite side.

## Introduction

The spontaneous assembly of proteins into liquid-like condensates plays an important role in numerous biological processes, from chromosomal organization and gene regulation to protein synthesis^1–4^. This phenomenon, which was initially observed for proteins residing in the cellular cytosol or in the nucleus, is now known to play a role in the assembly of diverse membrane-bound structures including the immunological synapse^5–7^, focal adhesions^8^, cell-cell junctions^9,10^, and endocytic vesicles^11^.

In each of these examples, proteins on both surfaces of the lipid bilayer are thought to play a role in condensate-driven assembly of biological structures. For example, during assembly of the immunological synapse, phase separation of linker for activation of T cells (LAT) on the cytoplasmic side of the membrane is likely to be reinforced by interactions among the extracellular domains of receptors^12^. Similarly, when focal adhesions are established, polymerization of integrin domains on the extracellular surface is complemented by phase separation of cytoplasmic proteins, such as talin, kindlin, and vinculin, on the intracellular side of the membrane^8^. Finally, during initiation of endocytic vesicles, phase separation of early endocytic proteins is likely complemented by condensation of transmembrane cargo proteins, such as receptor tyrosine kinases, several of which have recently been found to form liquid-like assemblies on membrane surfaces^13^. Collectively, these examples illustrate that proteins frequently phase separate simultaneously on both surfaces of lipid membranes. We sought to reconstitute this process in vitro so that we could directly observe and control the assembly of proteins on each surface of the membrane.

Conventional membrane substrates for in vitro reconstitution experiments include lipid vesicles^14,15^, which are closed spherical systems, and supported lipid bilayers^5,7,16–18^, in which a membrane is formed on a solid substrate. Protein phase separation has recently been observed on the outer surfaces of both types of membranes^15,16^. However, neither of these systems provides simultaneous access to both surfaces of the membrane. Therefore, we adapted a recently developed technique in which suspended planar lipid membranes are formed over the hexagonal holes of an electron microscopy grid^19,20^. Using this approach, we observed liquid-liquid phase separation (LLPS) simultaneously on both surfaces of a multiplexed planar membrane array.

Interestingly, we observed that phase separation of model proteins was highly coupled across the bilayer surface, such that protein-enriched regions on one side of the membrane tightly colocalized with protein-enriched regions on the opposite side of the membranes. Importantly, none of the proteins used in these experiments were transmembrane proteins or had the capacity for membrane penetration, such that there was no direct contact between proteins on opposite sides of the membrane surface. How then do protein-enriched domains communicate across the lipid membrane? Our results indicate that protein phase separation locally reduces lipid entropy, an effect which is minimized when protein-rich regions colocalize across the membrane barrier. These findings reveal a new fundamental mechanism of information transfer across lipid bilayer, which may play a role in initiating and stabilizing diverse cellular complexes that assemble at membrane surfaces.

## Results and Discussion

### The RGG domain of LAF-1 forms liquid-like condensates on freestanding planar membranes

Freestanding planar lipid membranes were created on the surface of the hexagonal holes of a transmission electron microscopy (TEM) grid, as described previously^19,20^ (Supplementary Fig. 1). Each grid contained 150 holes, each of which had a diameter of approximately 100 μm, enabling visualization of multiple independent membrane surfaces per field of view using fluorescence confocal microscopy.

To investigate protein phase separation on planar lipid membranes, the RGG domain of the LAF-1 protein (RGG) was selected because its participation in protein phase separation is well-established. Also, we chose RGG owing to its high solubility in aqueous buffers, such that phase separation in solution only occurs at RGG concentrations above several tens of μM^21^, about one to two orders of magnitude above the concentration used in our experiments with membranes. In this way, we could clearly differentiation protein phase separation on membranes from phase separation in the surrounding solution. We added RGG, labeled with Atto 488, with an N-terminal histidine-tag to freestanding planar membranes containing DGS-Ni-NTA lipids. Here, binding of his-RGG to the membrane is achieved through interactions between histidine and Ni-NTA (Fig. 1a). In our initial experiments, we sought to examine protein phase separation on only one surface of the bilayer. Therefore, we placed the TEM grid directly against a glass coverslip so that proteins only bound significantly to the top surface of the membrane.

**Figure 1.**
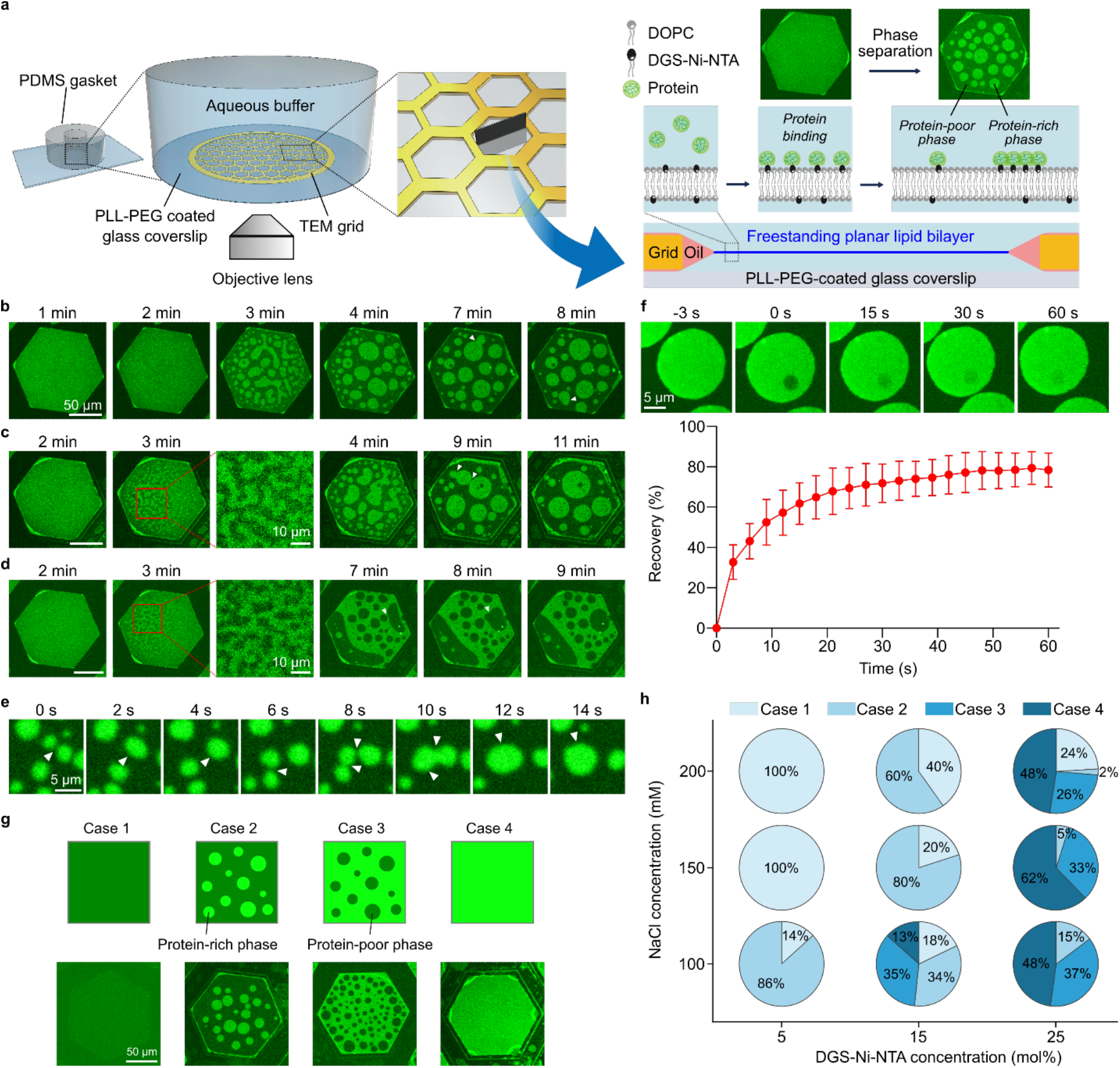
Phase separation of RGG domains on freestanding planar membranes results in liquid-like protein assemblies. **a**, Left: Schematic of the freestanding planar lipid membrane array system. Inside a PDMS chamber, a transmission electron microscopy (TEM) grid with hexagonal holes is immersed in an aqueous buffer, and freestanding planar membranes are created within multiple hexagonal holes. Right: Cross-section of the freestanding planar membrane spanning a single hexagonal hole. Once proteins (histidine-tagged) are added to the aqueous buffer in the chamber, they bind to the membrane through histidine-nickel interactions and phase separate into protein-rich and protein-poor phases. **b-d**, Representative images showing spinodal decomposition and the increase in domain size. The elapsed time since protein addition is indicated above each column. White arrowheads indicate fusion events. Scale bars, 50 µm. **e**, Fusion events between different protein-rich domains on the membrane over time. White arrowheads indicate fusion events. Scale bar, 5 µm. **f**, Top: Representative microscopic images showing fluorescence recovery after photobleaching (FRAP) of a protein-rich domain on the membrane. Bottom: Corresponding FRAP profile. Error bars indicate standard deviation (n = 3). Scale bar, 5 µm. **g**, Top: Schematic of four possible cases, from case 1 to 4, when phase-separating proteins are associated with membranes. Bottom: Representative microscopic images for each case. Scale bar, 50 µm. **h**, Percentage of individual cases as a function of DGS-Ni-NTA concentration in the membrane and NaCl concentration in the aqueous buffer. More than 130 lipid membranes were analyzed from 2-3 independent experiments for each condition. Membrane composition: 75 mol% DOPC, 25 mol% DGS-Ni-NTA (**b-d**), 85 mol% DOPC, 15 mol% DGS-Ni-NTA (**e**,**f**). 0.5 mol% Texas Red-DHPE was added for all. Buffer: 25 mM HEPES, 100 mM NaCl, pH 7.4. 1 µM of his-RGG labeled with Atto 488 were used.

Within minutes after adding the his-RGG protein, the protein began to bind to the top surface of the membrane. Initially, the fluorescence intensity in the protein channel was homogeneous over the surface of the membrane, indicating uniform protein binding. However, after a few minutes, heterogeneity in the intensity of the protein channel appeared, such that brighter and dimmer regions began to coexist, indicating phase separation of the membrane-bound protein layer into protein-rich (brighter) and protein-poor (dimmer) phases. We also observed that like phases, either protein-rich or protein-poor, fused and re-rounded upon contact, such that the phase separation coarsened over time (Fig. 1b-e, Supplementary Movie. 1,2). Interestingly, the protein-rich and protein-poor phases displayed a bicontinuous morphology at early moments after protein binding (Fig. 1b-d, 2-3 min). This observation suggests that phase separation occurred through spinodal decomposition, indicating that phase separation occurs spontaneously, owing to phase instability, rather than a nucleation and growth process^22–24^. In addition, protein-rich regions showed fluorescence recovery within a few seconds after photobleaching, with t_1/2_ of 10.5 s and mobile fraction of 79% (Fig. 1f). Collectively, these results suggest that two-dimensional, membrane-bound protein condensates of the RGG domain have liquid-like properties. Notably, because the membranes were composed of unsaturated lipids, DOPC and DGS-Ni-NTA, with melting temperatures far below room temperature, phase separation of the lipid membrane is unlikely to play a role in this process^25^. However, the impact of protein phase separation on the organization of the lipids is explored later in this report.

### The strength of protein-protein and protein-membrane interactions determine the area fraction of the protein-rich phase

We next probed the sensitivity of membrane-bound LLPS to changes in the strength of protein-protein and protein-membrane interactions. To vary the strength of protein-protein interactions, we changed the ionic strength of the buffer by controlling the sodium chloride (NaCl) concentration. Specifically, electrostatic attraction is known to play a major role in phase separation of RGG, such that increasing salt concentration screens residue-residue interactions, hindering phase separation^21^. To vary the strength of protein-membrane interactions, we changed the concentration of DGS-Ni-NTA lipid within the membrane.

Comparing the membranes within the grid upon addition of his-RGG, we observed four distinct cases, with the later cases becoming more common as protein concentration increased and ionic strength decreased (Fig. 1g). In the first case, the grid hole was covered completely by the protein-depleted phase. In the second case, the continuous phase was protein-depleted, while the dispersed phase was protein-enriched. In the third case, the continuous phase was protein-enriched, while the dispersed phase was protein-depleted. Finally, in the fourth case, the entire hole was covered by the protein-enriched phase. To characterize the impact of protein-protein and protein-lipid interactions on membrane-bound LLPS, we quantified the proportion of the grid holes that belonged to each of these cases, 15 minutes after protein addition. When LLPS was weakened by high ionic strength and low Ni-NTA concentration, the majority of grid holes belonged to cases 1 and 2. In contrast, when LLPS was strengthened by low ionic strength and high Ni-NTA concentration, the majority of the grid holes belonged to cases 3 and 4 (Fig. 1h). The observation of multiple cases for the same condition does not arise from variation in lipid composition between the grid holes, as a non-phase separating protein, green fluorescent protein (GFP) bound each hole approximately equally (Supplementary Fig. 2). However, it may arise from imperfect mixing upon addition of protein to solution and the cooperative relationship between protein binding and phase separation. Taken together, these results demonstrate that the same variables that control phase separation in solution – local protein concentration and the strength of protein-protein interaction – also govern protein phase separation on membrane surfaces.

### Increasing salt concentration lowers the critical temperature for phase separation of RGG on membrane surfaces

A characteristic of systems that undergo liquid-liquid phase separation is that the relative concentrations of macromolecules in the dilute and enriched phases become more similar to one another as the temperature of observation approaches the critical temperature^26^. Ultimately, when the critical temperature is reached, the two phases have the same concentration and therefore become indistinguishable from one another, such that one homogenous phase exists. To determine whether the membrane-bound condensates of the RGG domain display this behavior, we observed our system under conditions of increasing temperature. At each temperature, the relative fluorescence intensities of the protein-enriched and protein-depleted phases provide a rough estimate of the difference in protein concentration between the two phases^11^. In particular, by using these concentrations to represent the ends of a tie line, an approximate temperature-concentration phase diagram can be constructed (Fig. 2c), where C_rich_ is proportional to the protein concentration in the protein-enriched phase and C_poor_ is proportional to the protein concentration within the protein-depleted phase. We used this approach to map the phase diagram for membrane-bound condensates of RGG at an NaCl concentration of 100 mM. Here, the difference in intensity between the protein-enriched and protein-depleted regions was gradually lost as the temperature was raised from room temperature to 39 °C, which was the highest temperature we could achieve in our microscopy system. At 39 °C, some protein-enriched condensates of low contrast remained, suggesting that the critical temperature was greater than 39 °C at this NaCl concentration (Fig. 2a). In contrast, a complete dissolution of the protein-enriched phase was observed at 32 °C with an NaCl concentration of 200 mM (Fig. 2b). Our observation that the critical temperature decreases with increasing NaCl concentration is consistent with previous reports that increasing NaCl concentration screens residue-residue interactions between RGGs, hindering protein condensation^21^.

**Figure 2.**
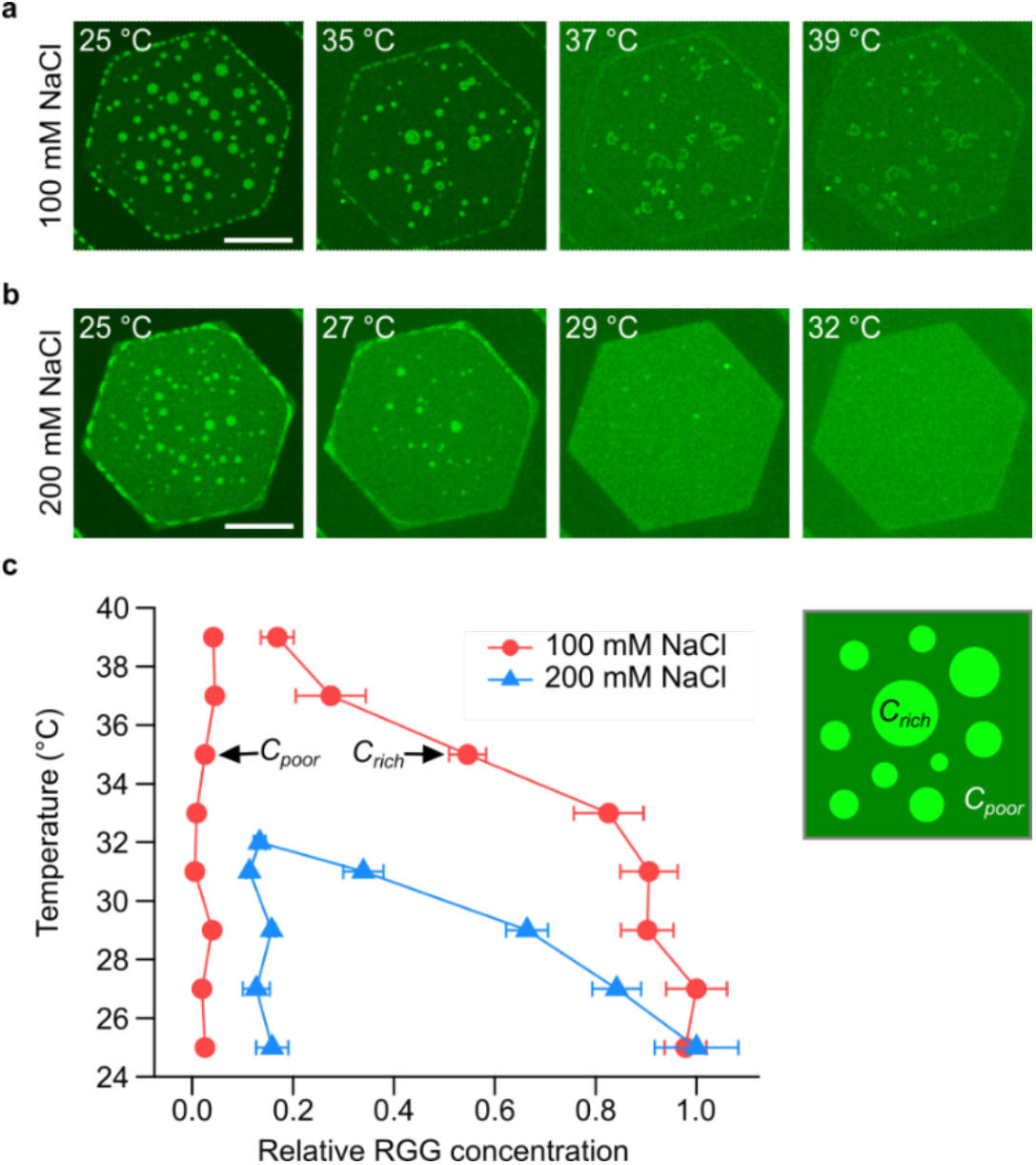
Concentration-temperature phase diagram with varying NaCl concentrations. **a**,**b**, Representative images at different temperature, indicated at the top left for each image, after addition of 1 µM of his-RGG, labeled with Atto 488, with the NaCl concentration of 100 mM **(a)** and 200 mM **(b)**. Membrane composition: 85 mol% DOPC, 15 mol% DGS-Ni-NTA, and 0.5 mol% Texas Red-DHPE. Scale bars, 50 µm. **c**, Phase diagram of RGG condensates on the membrane with varying temperature depending on NaCl concentration. *C*_*rich*_ and *C*_*poor*_ represent the concentration of protein-rich and protein-poor phases, respectively. Error bars represent standard deviations from analyzing multiple protein-rich and protein-poor regions (n > 10 for each) at each temperature.

### Transbilayer coupling of protein condensates occurs when proteins phase separate simultaneously on both sides of the membrane

So far, we have only allowed RGG proteins to bind and phase separate on one surface of the membrane. What will happen if phase separation occurs on both surfaces of the lipid bilayer simultaneously? By introducing thin spacers between the TEM grid and the glass coverslip at the bottom of our imaging chamber, both sides of the membrane were exposed to proteins dissolved in the surrounding aqueous medium (Fig. 3a). Within a few minutes after proteins bound to both sides of the membrane, we began to observe heterogeneity in the distribution of proteins over the membrane surface. Protein-enriched and protein-depleted regions appeared, as before. However, there was a key difference. When proteins bound to only one side of the membrane, we observed two levels of protein intensity, the brighter of which corresponded to the protein-enriched phase, while the dimmer corresponded to the protein-depleted phase (Fig. 1). In contrast, when proteins bound to both sides of the membrane, there were three levels of intensity, which we will denote as the brightest, medium-bright, and dimmest regions (Fig. 3b). Interestingly, we observed that the intensity difference between the dimmest and medium-bright regions was similar to that between medium-bright and the brightest regions. The simplest explanation for these observations is that the regions of dimmest intensity represent areas where proteins on both surfaces of the membrane are in the protein-depleted phase. In contrast, the regions of brightest intensity represent areas where proteins on both surfaces of the membrane are in the protein-enriched phase. Finally, the regions of medium intensity represent areas where proteins on one side of the membrane are in the protein-depleted phase and those on the other side of the membrane are in the protein-enriched phase (Fig. 3c). Importantly, the spacer used to support the suspended bilayer was thin, such that the bilayer remained within the limited working distance of the high magnification microscope objective. For this reason, the bottom surface of the suspended membrane was somewhat less accessible to proteins compared to the top surface. This difference resulted in slower protein binding and phase separation on the bottom surface, likely leading to smaller protein-enriched regions on the bottom surface compared to the top surface, particularly at early times (Fig. 3c cartoon). Interestingly, regions of brightest intensity were typically surrounded by regions of medium intensity. This observation suggests that protein phase separation on one side of the membrane was coupled to protein phase separation on the other side of the membrane, such that phase-separated regions tended to colocalize across the membrane boundary.

**Figure 3.**
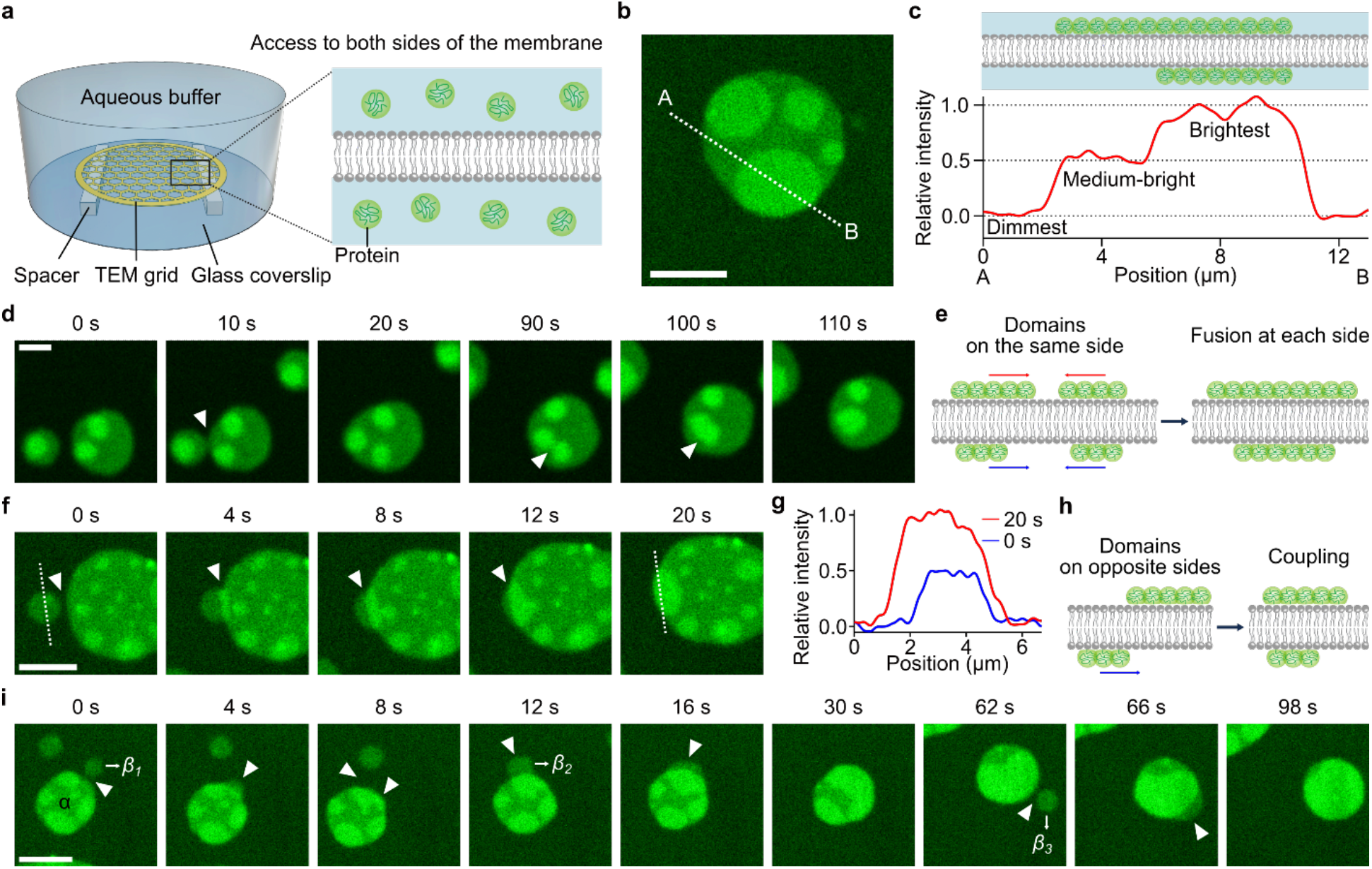
Simultaneous protein phase separation on both membrane surfaces leads to transbilayer coupling of protein-rich domains. **a**, Schematic of the system where protein access to both sides of the membrane is possible by placing spacers between the TEM grid and the glass coverslip. **b**, Representative microscopic images of protein regions with three different brightnesses: dimmest, medium-bright, and brightest regions. **c**, Cartoon of cross-section of the membrane (Top) and relative intensity profile (Bottom) along the dotted line (from A to B) in **b**, where regions with relative intensity of 0, 0.5, and 1.0 correspond to the dimmest, medium-bright, and brightest regions, respectively. Relative intensity (*I*_*R*_) was defined as *I*_*R*_ *= (I – I*_*D*_*)/(I*_*B*_ *– I*_*D*_), where *I, I*_*D*_, and *I*_*B*_, indicate the fluorescence intensity of the region of interest, the intensity of the dimmest region, and the intensity of the brightest region, respectively. **d-i**, Representative microscopic images over time and cartoons showing dynamic changes when different protein domains were spatially overlapping. Fusion occurred when domains were on the same side of the membrane (**d**,**e**), and coupling occurred when domains were on opposite sides of the membrane (**f-i**). **g**, Relative intensity profile along the dotted white lines in the first (0 s, blue) and the last (20 s, red) images in **f**. White arrowheads indicate fusion (**d**) or coupling (**f**,**i**) spots. 1 µM of his-RGG labeled with Atto 488 was used. Buffer: 25 mM HEPES, 100 mM NaCl, pH 7.4. Membrane composition: 85 mol% DOPC, 15 mol% DGS-Ni-NTA, and 0.5 mol% Texas Red-DHPE. Scale bars, 5 µm.

If protein phase separation is coupled across the membrane bilayer, what should happen when two protein-enriched domains cross paths? If they are on the same side of the membrane, they should fuse together upon contact. In line with this prediction, Figure 3d shows two medium-bright domains that each contain one or two domains of the brightest intensity. When the medium-bright domains contacted one another, they fused together within seconds, similar to our observation of fusion between domains on a single side of the membrane (Fig. 1b-e). This fusion event brought the domains of brightest intensity into contact with one another, also resulting in their fusion (Fig. 3d, 90 s, Supplementary Movie. 3). These observations demonstrate that protein-rich domains on the same side of the membrane fuse together upon contact (Fig. 3e), which provides further evidence that protein phase separation is occurring on both surfaces of the membrane.

Next, if coupling is occurring, when two protein-enriched domains are on opposite sides of the membrane and cross paths, they should become coupled. In line with this prediction, Figure 3f (Supplementary Movie. 4) shows a medium-bright region that meets another, larger medium-bright region, which also includes several smaller regions of brightest intensity. As the two medium-bright regions meet, rather than fusing, a stable overlap region develops, which has a brightness similar to the brightest regions within the same image, showing a stepwise increase in relative intensity from 0.5 to 1.0 (Fig. 3g). Based on our previous observations, if these two domains were on the same side of the membrane, then fusion between them would have occurred with the brightness unchanged and the overall domain size increasing. Instead, the observed increase in brightness, without a change in domain size, strongly suggests that medium-bright domains on opposite sides of the membrane crossed paths, recognized each other, and became coupled (Fig. 3h). Similarly, we observed consecutive domain coupling events, which eventually led to full transmembrane coupling of protein-enriched domains (Fig. 3i, Supplementary Movie. 5). Specifically, we initially observed a protein region, denoted as the *α* region, composed of both medium-bright domains and domains of brightest intensity. Then, between 0 s and 8 s, the medium-bright part of the *α* region and another smaller medium-bright region, denoted as the *β*_*1*_, crossed paths. Since brightness increased in the overlapped region without an increase in domain size, it appeared that these two overlapping domains were on different sides of the membrane and became coupled. After that, two more similar coupling events were observed (12 s – 16 s and 62 s – 66 s), as the *β*_*2*_ and *β*_*3*_ domains, initially having medium brightness, became coupled, such that full coupling was achieved, resulting in a single domain of brightest intensity (Fig. 3i).

In all cases, protein-enriched domains either merged, if on the same side of the membrane, or became stably coupled, if on opposite sides of the membrane. In contrast, if protein-enriched regions on opposite sides of the membrane were uncoupled, we would expect them to diffuse over one another without becoming coupled. Such an event would transiently create a region of brightest intensity during the time that the domains passed over one another on opposite sides of the membrane, but would not lead to stable coupling. Events with these characteristics were never observed in our experiments. Collectively, our observations demonstrate that protein-enriched regions on the surfaces of suspended lipid bilayers undergo transmembrane coupling, such that protein phase separation on one side of the membrane frequently colocalizes with protein phase separation on the opposite side of the membrane. How can protein condensates on different sides of the membrane recognize each other and become coupled? Because the proteins attach peripherally to the membrane surface via the histidine-Ni-NTA interaction, there is no direct contact between proteins on the two sides of the membrane, suggesting that coupling occurs indirectly through protein-lipid interactions. Therefore, we next examined the impact of protein condensates on the behavior of the membrane lipids.

### Lipid probes are depleted from protein-rich regions, suggesting that protein phase separation locally orders membrane lipids

To probe the impact of protein phase separation on the lipids, we began by examining the distribution of a fluorescent lipid probe, Texas Red-DHPE, which we included at 0.5 mol% in the solvent mixture used to create the suspended membranes. Interestingly, the intensity distribution in the lipid channel was opposite of that in the protein channel, such that the brightest regions in the protein channel, which are the coupled protein-enriched regions, corresponded to the dimmest regions in the lipid channel. Likewise, the dimmest regions in the protein channel, which are the protein-depleted regions, corresponded to the brightest regions in the lipid channel (Fig. 4a). These observations suggest that protein phase separation results in the depletion of the probe lipids from the underlying membrane. In particular, in the lipid probe channel, the increase in fluorescence intensity from the dimmest to medium-bright regions was comparable to the increase from the medium-bright to the brightest regions (Fig. 4b). This comparison suggests that protein condensation on both sides of the membrane resulted in about twice as much depletion of the probe lipid as protein condensation on one side of the membrane. Here, to check if a small amount of residual oil (hexadecane) in our membrane had an effect on lipid probe exclusion, we replaced hexadecane (C_16_H_34_) with squalane (C_30_H_62_), which has a longer hydrocarbon chain that has been shown to greatly reduce the amount of trapped oil in the bilayer, leading to essentially solvent-free membranes^27–29^. We observed a very similar depletion of the probe lipid from the protein-enriched regions using squalane (Supplementary Fig. 3), suggesting that depletion of the probe lipid cannot be explained by inclusion of oil in the bilayer.

**Figure 4.**
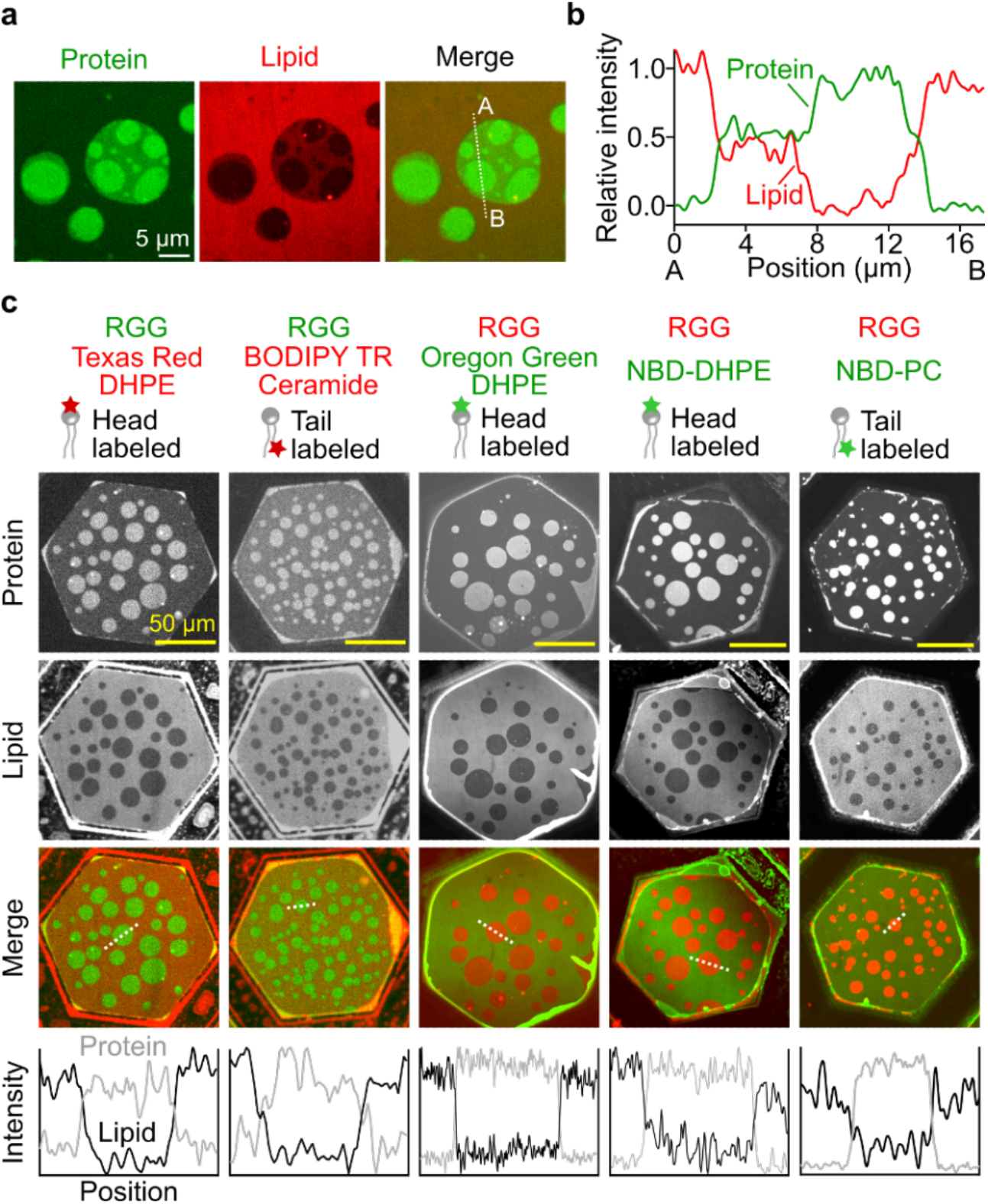
Multiple lipid probes partition away from protein-rich regions. **a**, Representative microscopic images showing regions with three different brightness from protein and lipid channels. 1 µM of his-RGG labeled with Atto 488 was applied to both sides of the membrane. Membrane composition: 85 mol% DOPC, 15 mol% DGS-Ni-NTA, and 0.5 mol% Texas Red-DHPE. Scale bar, 5 µm. **b**, Relative intensity profile along the dotted line (from A to B) in the merged channel in **a**, where green and red lines indicate relative intensity from protein and lipid channels, respectively. See caption of Figure 3c for the definition of relative intensity. **c**, Representative microscopic images after the addition of 1 µM of his-RGG to the top side only of each membrane containing either head labeled (Texas Red-DHPE, Oregon Green-DHPE, and NBD-DHPE) or tail labeled (BODIPY TR-Ceramide and NBD-PC) lipid probes. His-RGG was labeled with Atto 488 or Atto 594 depending on lipid probes used. Fluorescence intensity profile along the dotted line in each merged channel image is shown at the bottom, where gray and black lines represent the intensity from protein and lipid channels, respectively. Buffer: 25 mM HEPES, 100 mM NaCl, pH 7.4. Membrane composition: 85 mol% DOPC, 15 mol% DGS-Ni-NTA, and 0.5-1.0 mol% lipid probe. Scale bars, 50 µm.

Next, we tested the generality of lipid probe exclusion from protein-enriched phases by examining the partitioning of several additional probe lipids in membranes on which protein phase separation was taking place. Texas Red-DHPE emits in the orange/red region of the spectrum, and is covalently conjugated to the head group of a phospholipid. In addition to this probe, we also examined Oregon Green-DHPE and NBD-DHPE, which are similar to Texas Red-DHPE in that the fluorophore is conjugated to the phospholipid head group, except that the incorporated fluorophores emit in the green region of the spectrum. We also examined BODIPY TR-Ceramide (orange/red) and NBD-PC (green), which incorporate fluorophores conjugated to the lipid tail group, eliminating direct interactions between proteins and fluorescent probes. We examined the partitioning of each of these probe lipids in membranes on which phase separation of RGG was taking place. For simplicity, we only looked at the case in which membrane binding and phase separation of RGG was restricted to one side of the membrane. In each case we observed that the dimmer regions in the lipid channel corresponded to the protein-enriched regions that were brighter in the protein channel (Fig. 4c). These data illustrate that a diverse set of probe lipids are depleted from regions of the membrane on which protein-enriched phases exist. We also confirmed that lipid probe partitioning was observed when unlabeled his-RGG was used (Supplementary Fig. 4), demonstrating that the reduction in probe lipid intensity in protein-rich regions could not be explained by spectral interactions between their respective fluorophores.

Why are diverse probe lipids excluded from the protein-enriched phase? Notably, the conjugation of a fluorophore to a lipid substantially increases its molecular weight, creating a bulky amphiphile that often disrupts the packing of other membrane lipids. As a result, most probe lipids are known to be excluded from liquid-ordered and solid-like membrane phases^30,31^. This reasoning suggests that assembly of protein-rich condensates on membrane surfaces locally orders lipids in the underlying membrane, resulting in reduced localization of probe lipids to these regions. Notably, the term ordered lipids does not refer to phase-separated lipid regions composed of saturated lipids and sterols, as reported elsewhere in the literature^30,31^. Instead, we use the term ‘lipid ordering’ to indicate that the conformational freedom and entropy of the lipids are locally reduced due to the presence of protein condensates.

### Transmembrane association of protein condensates is consistent with an entropic coupling mechanism

Results in the previous section suggest that assembly of protein condensates on the surface of a membrane locally orders the underlying lipids. How might local ordering of lipids contribute to transmembrane association of protein condensates? Local ordering of lipids implies a reduction in thermal fluctuations among the lipids. Previous work has suggested that when one membrane leaflet becomes ordered, fluctuations in the opposing membrane leaflet are also suppressed^32,33^, which reduces the system’s entropy. The resulting reduction in entropy can be minimized by transmembrane association of regions with ordered lipids, thereby maximizing fluctuations among the surrounding disordered portions of the membrane (Fig. 5a). This entropic coupling hypothesis makes several testable predictions. First, regions of the membrane with coupled protein condensates on both sides of the membrane should be less fluid in comparison to uncoupled regions with a protein condensate on only one side of the membrane. For this reason, we would expect coupled regions to fuse together more slowly than uncoupled regions. To evaluate this prediction, we monitored, in the lipid channel, fusion between uncoupled (medium-bright) and coupled (dimmest) regions (Fig. 5b,c). We quantified the relaxation after fusion by measuring the aspect ratio of two coalescing domains over time, which followed an exponential decay^34^. We observed that coupled regions had a longer characteristic relaxation time than uncoupled ones (Fig. 5d), which suggests that protein phase separation reduced the fluidity of the resulting protein-membrane composite.

**Figure 5.**
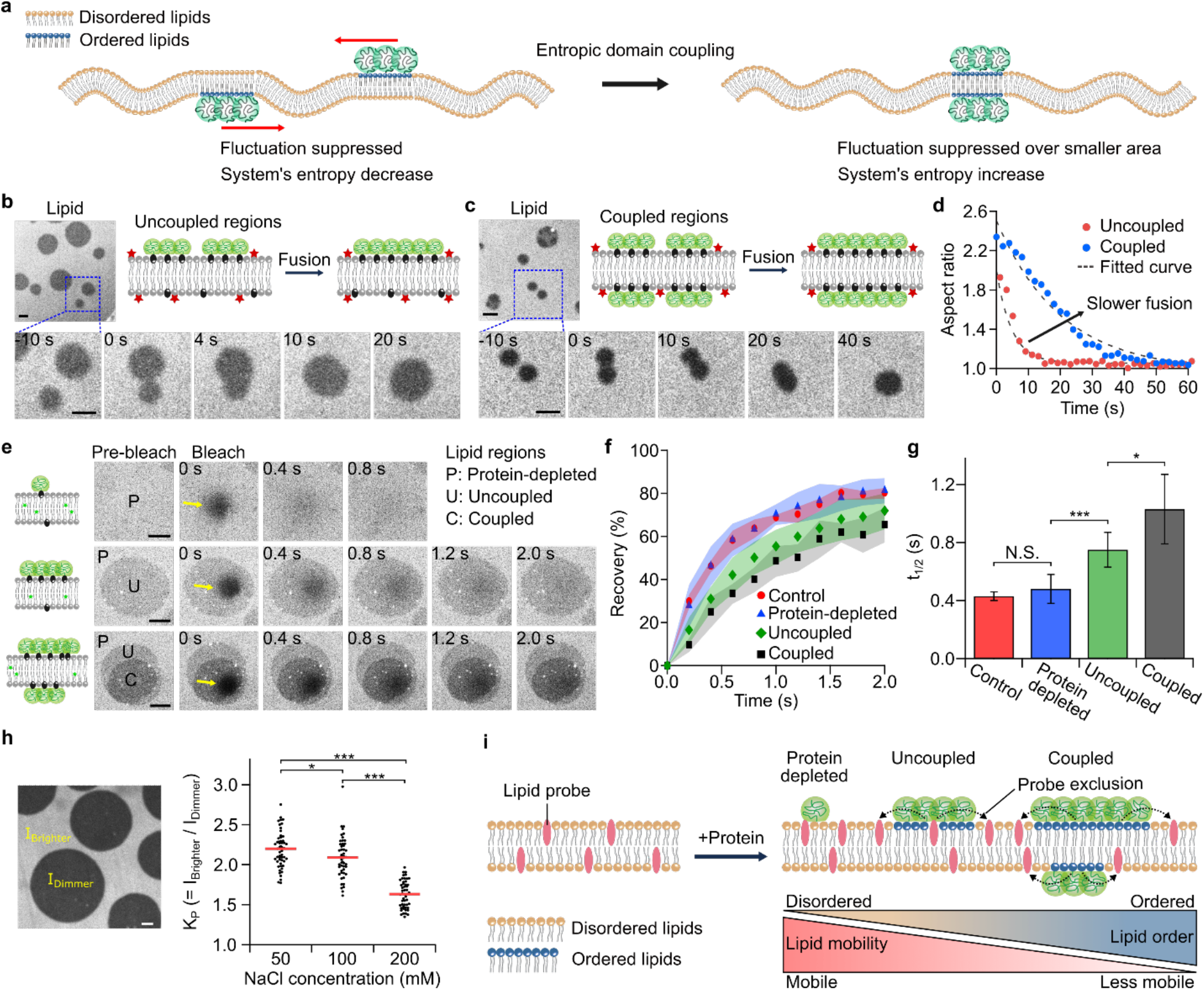
Protein assemblies on membranes create ordered lipid regions. **a**, Schematic of the entropic coupling mechanism. Left: Protein condensates induce ordered lipid regions (lipids with blue heads) and thermal fluctuations of these regions become suppressed. Right: Protein condensates on different sides of the membrane become coupled, minimizing free energy. **b**,**c**, Representative microscopic images from lipid channels and cartoons showing fusion events of two uncoupled regions (**b**) and two coupled regions (**c**) of membrane-protein composites over time. **d**, Aspect ratio changes over time during relaxation after fusion of two regions. Red circles indicate aspect ratio change for uncoupled regions, and blue circles for coupled regions. Dotted lines represent an exponential fit: y(t) = A + B*exp(-t/τ). **e**, Representative microscopic images in the lipid channel showing fluorescence recovery for protein-depleted, uncoupled, and coupled regions. Yellow arrows indicate photobleached regions. **f**, FRAP profile for the control (protein-free membrane, red circles), Protein-depleted (blue triangles), uncoupled (green diamonds), and coupled (black squares) regions. Shaded regions in each color represent standard deviations for each case from 5 to 17 independent measurements. **g**, Corresponding t_1/2_, time required for 50% of fluorescence recovery, from FRAP profile in **f**. Error bars indicate standard deviation. **h**, Left: Representative microscopic image in the lipid channel showing both brighter and dimmer regions for calculating partition coefficient of lipid probe. Right: Partition coefficients (*KP*) as a function of different NaCl concentrations. The red bars indicate the average *KP* values for each case. **i**, Schematic of hypothesis that protein condensates induce ordered lipid regions with reduced lipid mobility. Brackets in **g** and **h** show statistically significant comparisons using an unpaired, two-tailed Student’s t test. *p < 0.05, ***p < 0.001, and N.S. indicates a difference that was not statistically significant. Membrane composition: 85 mol% DOPC, 15 mol% DGS-Ni-NTA with 0.5 mol% Texas Red-DHPE (**b**,**c**,**h**) or NBD-PC (**e**).1 µM of unlabeled his-RGG was used. Scale bars, 5 µm.

If protein condensation reduces the fluidity of the membrane, then the lipids beneath the condensates should experience slower diffusion in comparison to lipids in regions with more dilute protein binding. To test this second prediction, we used fluorescence recovery after photobleaching (FRAP) to examine the diffusion of probe lipids within: (i) protein-depleted regions (brightest in the probe lipid channel), (ii) regions with uncoupled protein-enriched domain on a single side of the bilayer (medium-bright regions in the probe lipid channel), and (iii) regions with coupled protein-enriched domains on both sides of the bilayer (dimmest regions in the probe lipid channel) (Fig. 5e). Here we used the NBD-PC probe lipid, which is tail-labeled, such that the NBD fluorophore is unlikely to interact with the protein layer. Unlabeled RGG proteins were used to eliminate the possibility of spectral cross-talk with the probe lipid. The time required for 50% recovery after bleaching, *t*_*1/2*_, was obtained from the recovery curve for each region (Fig. 5f). We observed that the brightest lipid region exhibited *t*_*1/2*_ of 0.48 s, similar to that of protein-free control membranes (0.43 s), whereas medium-bright and dimmest lipid regions displayed longer *t*_*1/2*_ of 0.75 s and 1.03 s, respectively (Fig. 5g). Slower recovery suggests that lipid mobility within the region of interest was reduced. From these results we conclude that the diffusivity of the probe lipid was greatest in the protein-depleted phase, decreased in regions of the membrane with protein-enriched phase on one side, and decreased further in regions of the membrane with coupled protein-enriched phases on both sides.

Collectively, these results support the idea that protein condensation reduces the entropy of the underlying lipids. If protein phase separation is responsible for ordering the membrane and depleting probe lipids from protein-enriched phases, then we would expect that strengthening protein-protein interactions would increase the extent of probe lipid depletion. To test this final prediction, we defined a partition coefficient (*K*_*P*_) as *K*_*P*_ *= I*_*B*_ */ I*_*D*_, where *I*_*B*_ and *I*_*D*_ indicate the fluorescence intensity of the brighter and the dimmer regions in the lipid channel, respectively^35^. Measurements of *K*_*P*_ were conducted for the case in which protein phase separation took place on only one side of the membrane after applying 1 μM of his-RGG, where Texas Red-DHPE was used as a lipid probe. Based on the definition of *K*_*P*_, the more the probes are excluded from the dimmer region, the higher the *K*_*P*_ value will become. We observed that decreasing the concentration of NaCl, which is expected to strengthen interactions among RGG domains, resulted in increasing values of *K*_*P*_ (Fig. 5h). This result demonstrates that when protein-protein interactions are strengthened, the membrane becomes more ordered, resulting in greater depletion of probe lipids from the underlying lipid bilayer.

### Domain coupling is a general phenomenon when proteins phase separate on membrane surfaces

Collectively, the results in the previous section support the hypothesis that entropic coupling drives transmembrane colocalization of protein phase separation. However, so far we have only examined phase separation of the RGG domain. To examine the generality of this mechanism, we examined membrane-bound phase separation of an additional protein domain known to form liquid-like condensates, the low complexity domain of fused in sarcoma (FUS LC)^36^. Similar to our experiments with RGG, a histidine-Ni-NTA interaction was used to attract the protein on the membrane surface. We observed liquid-like assemblies of FUS LC on membranes, which fused and re-rounded upon contact. Additionally, we observed a similar trend of probe lipid partitioning, where lipid probes were depleted from protein-enriched regions (Fig. 6a). We also observed transbilayer domain coupling when FUS LC proteins were allowed to phase separate on both sides of the membrane simultaneously (Fig. 6b,c). Importantly, phase separation of FUS LC relies mainly on pi-pi stacking interactions among tyrosine residues, whereas phase separation of RGG is dominated by electrostatic interactions. Given these differences, the very similar impact of the two proteins on membrane organization suggests that the transbilayer coupling phenomenon observed here is a general mechanism that may be applicable to diverse proteins that phase separate at membrane surfaces.

**Figure 6.**
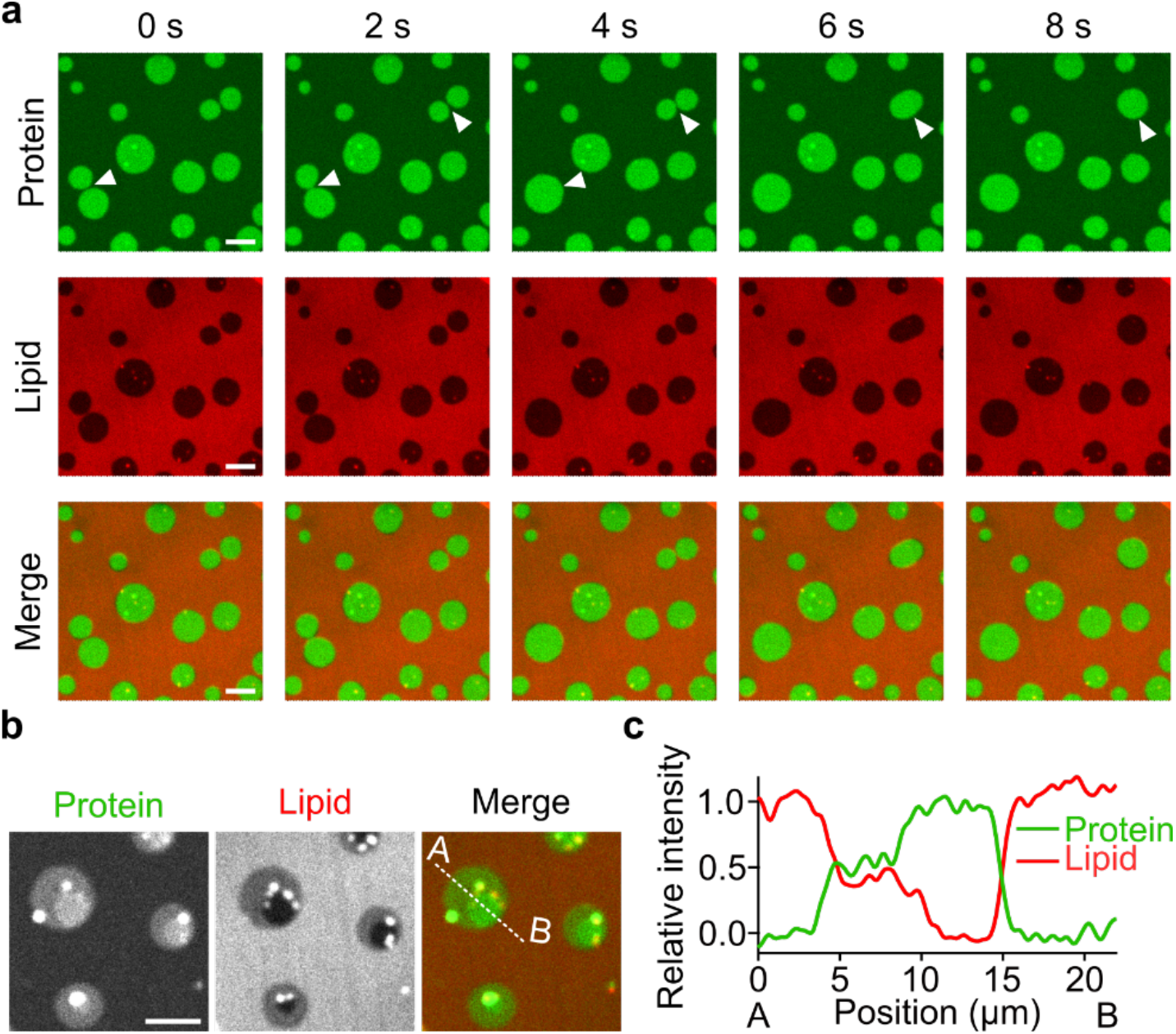
Phase separation of FUS LC domains on membranes. **a**, Representative microscopic images showing fusion of protein-rich regions. White arrowheads indicate fusion events. **b**, Representative microscopic images showing transbilayer domain coupling. Scale bars, 5 µm. **c**, Relative intensity profiles along the dotted lines (from A to B) in the merged channel in **b**, where green and red lines represent the intensity from protein and lipid channels, respectively. See caption of Figure 3c for the definition of relative intensity. 500 nM of his-FUS LC, labeled with Atto 488, was used. Membrane composition: 80 mol% DOPC, 20 mol% DGS-Ni-NTA, and 0.5 mol% Texas Red-DHPE. Buffer: 25 mM HEPES, 150 mM NaCl, pH 7.4. Scale bars, 10 µm.

## Discussion

Molecular reconstitution has provided fundamental insights into the mechanisms behind protein phase separation, both in solution^1–4^, and at membrane surfaces^5,7,14–18^. However, because existing membrane substrates for *in vitro* reconstitution provide access to only one side of the membrane, interactions between phase separated regions on opposite sides of a membrane surface have not been investigated.

Here, we introduce a freestanding planar membrane array as an appropriate platform to study membrane-associated protein phase separation simultaneously on both sides of a membrane surface. Using this approach, our images reveal coupling between protein-enriched condensates on one side of the membrane with those on the opposite side. In particular, our findings suggest that liquid-like protein assemblies on membranes, regardless of the identity of proteins, create ordered lipid regions with reduced lipid mobility, which become entropically coupled to ordered regions within the opposite leaflet. Notably, transmembrane coupling in our system was highly stable, such that, once they became coupled, membrane-bound protein condensates on opposite sides of the membrane were never observed to separate. The reduction in free energy owing to transmembrane coupling of liquid-ordered lipid domains has been estimated to be approximately 0.016 *k*_*B*_*T*/nm^2^ ^37^. Given that the coupled domains in our experiments have micrometer dimensions, the energetic barrier to uncoupling would be on the order of ∼ 10^4^ *k*_*B*_*T*, suggesting highly stable coupling. By the same logic, a coupled domain of 20-30 nanometers in diameter should incur a barrier to uncoupling of approximately 10 *k*_*B*_*T*, significantly above the thermal energy. These arguments suggest that stable coupling could extend to small length scales, relevant to many physiologically important structures. However, the precise scaling between domain size and stability remains to be measured.

From a biophysical perspective, the coupling mechanism identified in this work constitutes a new way of transferring information across biological membranes, which is independent of membrane-spanning proteins and lipid-lipid immiscibility. In particular, we demonstrate that a protein condensate on one side of the membrane can be detected by a condensate on the other side of the membrane through their mutual influence on the conformational freedom of the underlying lipids, a process that does not require a discontinuity in lipid composition, or direct contact between proteins on the two surfaces of the membrane.

From a biological perspective, it is increasingly clear that liquid-like protein condensates help to organize critical structures and events at biological membranes, from assembly of cell-cell junctions to the budding of trafficking vesicles^9–11^. Importantly, each of these assemblies involves protein-protein interactions on both surfaces of the membrane. In such processes, we speculate that the entropic coupling mechanism identified here works in concert with transmembrane proteins and lipid phase separation^38^ to achieve robust transbilayer coupling and communication.

## Acknowledgements

This research was supported by the NIH through grant R35GM139531 (J.C.S), the NSF DMS through grant 1934411 (J.C.S), and the Welch Foundation through grant F-2047 (J.C.S).

