## Supplementary Information for "Lateral compression of lipids drives transbilayer coupling of liquid-like protein condensates"

### Experimental Sections

#### Materials

1,2-dioleoyl-sn-glycero-3-phosphocholine (DOPC), 1,2-dioleoyl-sn-glycero-3-[(N-(5-amino-1-carboxypentyl)iminodiacetic acid)succinyl] nickel salt (DGS-Ni-NTA), and 1-palmitoyl-2-{6-[(7-nitro-2-1,3-benzoxadiazol-4-yl)amino]hexanoyl}-sn-glycero-3-phosphocholine (NBD-PC) were purchased from Avanti Polar Lipids. Texas Red 1,2-dihexadecanoyl-sn-glycero-3-phosphoethanolamine triethylammonium salt (TR-DHPE), BODIPY TR Ceramide (TR-Ceramide), Oregon Green 488 1,2-dihexadecanoyl-sn-glycero-3-phosphoethanolamine (OG-DHPE), N-(7-nitrobenz-2-oxa-1,3-diazol-4-yl)-1,2-dihexadecanoyl-sn-glycero-3-phosphoethanolamine triethylammonium salt (NBD-DHPE), and 4-(2-hydroxymethyl)-1-piparazineethanesulfonic acid (HEPES) were purchased from Thermo Fisher Scientific. Sodium chloride, sodium tetraborate, hexadecane, squalane, silicone oil AR 20, poly-L-lysine MW 15,000-30,000 (PLL), Atto 488 NHS ester, and Atto 594 NHS ester were purchased from Sigma-Aldrich. Amine-reactive PEG (mPEG–succinimidyl valerate, MW 5000) was purchased from Laysan Bio.

#### Protein expression and purification

Expression and purification of the RGG domain of LAF-1 protein (RGG) was performed as described previously[1]. Briefly, *E. Coli* BL21(DE3) competent cells were transformed with a plasmid encoding the RGG domain of LAF-1. After transformation, cells were grown in 1 liter of 2xYT media for 3-4 hours at 37 °C while shaking at 220 rpm until the optical density at 600 nm (OD 600) of the media became reached 0.8, followed by overnight expression induced with 0.5 mM of isopropyl β-D-1-thiogalactopyranoside (IPTG) at 18 °C while shaking at 220 rpm. Pellets of cells expressing RGG were harvested through centrifugation at 4 °C. Pellets were resuspended in 40 mL buffer containing 20 mM Tris, 500mM NaCl, 20 mM imidazole, 1% Triton X-100, and one EDTA-free protease inhibitor tablet (Sigma Aldrich) at pH 7.5, and lysed by sonication on ice. To prevent formation of RGG condensates, all of the following steps were done at room temperature. The cell lysate was clarified by centrifugation at 15,000 × g for 30 min and then incubated with Ni-NTA resin (G Biosciences, USA) for 1 hour to achieve binding. Protein-bound Ni-NTA resin was settled in a glass column and washed with a buffer containing 20 mM Tris, 500 mM NaCl, 20 mM imidazole, at pH 7.5. The bound proteins were eluted from the Ni-NTA resin with a buffer containing 20 mM Tris, 500 mM NaCl, 500 mM imidazole, at pH 7.5. Purified proteins were then buffer exchanged into the storage buffer (20 mM Tris, 500 mM NaCl, pH 7.5). Small aliquots of the protein were flash frozen using liquid nitrogen and stored at - 80 °C.

Expression and purification of the low-complexity domain of fused in sarcoma (FUS LC) was carried out according to previous reports[2,3]. In brief, FUS LC was overexpressed in *E. Coli* BL21(DE3) cells. Cells were grown for 4 hours at 37 °C while shaking at 220 rpm until OD 600 reached 0.8, followed by expression induced with 1 mM of IPTG at 37 °C for 3 hours. Pellets of cells expressing FUS LC were collected and lysed by sonicating on ice in 40 mL lysis buffer containing 500 mM Tris pH 8.0, 5 mM EDTA, 5% glycerol, 10 mM β-mercaptoethanol, 1 mM phenylmethylsulfonyl fluoride (PMSF), 1% Triton X-100 and one EDTA-free protease inhibitor tablet (Sigma Aldrich). The cell lysates were centrifuged at 40,000 rpm for 40 min at 4°C. Unlike RGG, FUS LC resided in the insoluble fraction after centrifugation. Therefore, the insoluble fraction was kept and resuspended in 8 M urea, 20 mM NaPi pH 7.4, 300 mM NaCl and 10 mM imidazole. The resuspended sample was then centrifuged at 40,000 rpm for 40 min at 4°C. Under denaturing conditions, FUS LC is soluble and resided in the supernatant. This supernatant was then mixed with Ni-NTA resin (G Biosciences, USA) for 1 hour at 4°C. The Ni-NTA resin was then settled in a glass column and washed with the above solubilizing buffer. The bound proteins were eluted from the Ni-NTA resin with a buffer containing 8 M urea, 20 mM NaPi pH 7.4, 300 mM NaCl and 500 mM imidazole. Purified proteins were then buffer exchanged into storage buffer (20 mM CAPS, pH 11) using 3K Amicon Ultra centrifugal filters (Millipore, USA). Small aliquots of the protein were flash frozen using liquid nitrogen and stored at - 80 °C.

### Protein labelling

RGG and FUS LC were labeled with the amine-reactive Atto 488 (or Atto 594) NHS ester for visualization. The labeling reaction for RGG took place in its storage buffer (25 mM HEPES, 500 mM NaCl, pH 7.5). Dye was added to the protein in 2-fold stoichiometric excess and allowed to react for 30 min at room temperature. Labeled protein was separated from unconjugated dye using 3K Amicon column. The labeling ratio was measured using UV–Vis spectroscopy. Labeled proteins were dispensed into small aliquots, flash frozen in liquid nitrogen and stored at - 80 °C. Labeling of FUS LC followed the same process except that the labeling reaction was done in the buffer containing 50 mM HEPES at pH 7.4, and the labeled protein then exchanged back into the storage buffer (20 mM CAPS, pH 11), divided into small aliquots, flash frozen in liquid nitrogen and stored at - 80 °C.

### Freestanding planar lipid membrane formation

Freestanding planar lipid membranes were prepared as described previously[4,5] with slight modifications (Supplementary Fig. 1). First, lipids dissolved in chloroform were mixed in a glass vial and dried under a gentle $N_2$ stream. The dried lipid film was redissolved in a mixture of hexadecane and silicone oil (1:1 v/v) to obtain a lipid/oil solution with a total lipid concentration of 3 mM. Each oil was filtered through 0.2 µm syringe filter (Corning Inc.) before use. The lipid/oil solution was bath-sonicated for 30 min and used for experiments within several hours.

The imaging chamber was assembled by placing a PDMS gaskets onto no. 1.5 glass coverslips (VWR). To make a PDMS gasket, 2.1 g of Sylgard™ 184 silicone elastomer base and 0.3 g of curing agent (Dow Corning) were thoroughly mixed and poured into a single well of a 12-well plate, where a cylindrical structure with a diameter of 8 mm was placed at the center to make a circular hole in the PDMS layer. The mixture was then incubated at 45 °C for at least 3 hours for curing. Coverslips were cleaned with 2% v/v Hellmanex III (Hellma Analytics) solution, rinsed thoroughly with deionized water, and dried under a gentle $N_2$ stream prior to use. The coverslip was then passivated with a layer of PLL-PEG, which was synthesized as described previously[6]. In each imaging chamber, 15 µL of PLL-PEG solution was added, followed by repetitive rinsing with the experimental buffer using a pipette after 20 min of incubation. A total of 150 µL of the aqueous buffer was then added into the PLL-PEG passivated imaging chamber. After that, 2 µL of lipid/oil solution was gently dropped and spread on the air-aqueous buffer interface. After several minutes, a hexagonal TEM grid made of gold (G150HEX Au, Gilder Grids), which was hydrophobically coated by thiol-gold reactions before use, was gently placed on the air-oil interface using tweezers. The grid was placed there for several minutes so that a thin oil film could be created within the grid holes. Then, the grid was submerged into the aqueous buffer using a syringe needle to place it on the PLL-PEG coated glass surface. The thickness of the oil film decreased as the oil drained out, and spontaneous adhesion of two lipid monolayers occurred, resulting in a freestanding planar lipid bilayer. Proteins were then added to the aqueous buffer above the lipid bilayer.

In experiments requiring a spacer, two small strips of Scotch® Magic™ Tape (3M), each a few millimeters wide were adhered to the coverslip surface. The space between these strips was a few millimeters, slightly less than the diameter of the grid. To form the membranes, the same procedure was followed to coat the grid with the lipid oil solution. When the grid was submerged in the aqueous buffer, care was taken to settle it on top of the strips of tape, so that it bridged the gap between them, such that the protein solution could flow beneath the grid.

### Microscopy

A spinning disc confocal microscope (SpinSR10, Olympus) equipped with a Hamamatsu Orca Flash 4.0V3 sCMOS Digital Camera was used to visualize samples. 20X objective (1-U2M345, Olympus), 1.40 NA/40X oil immersion objective (1-UXB220, Olympus), and 1.50 NA/100X oil immersion objective (1-UXB170, Olympus) were used for visualization. Laser wavelengths of 488 and 561 nm were used for excitation. FRAP was performed using the Olympus FRAP unit 405 nm laser and 1.50 NA/100X oil immersion objective (1-UXB170, Olympus).

### Image analysis

ImageJ was used for image analysis. For time-lapse image sets, contrast and brightness were kept the same for all images. For all cases, fluorescence intensity values were measured in unprocessed images.

For FRAP analysis, the FRAP Profiler plugin for ImageJ was used. Fluorescence recovery of regions with 2-5 μm diameter were measured over time, and intensity values were normalized to the maximum pre-bleach and minimum post-bleach values.

For the fusion dynamics of two coalescing membrane-protein composite regions, observed in the lipid channel, aspect ratio values were measured using the “Analyze Particles” function.

For measurement of partition coefficients ($K_P$) of Texas Red DHPE under different NaCl concentrations, fluorescence intensity values (background subtracted) of two adjacent brighter and dimmer regions were used to avoid the effect of locally variations in intensity.

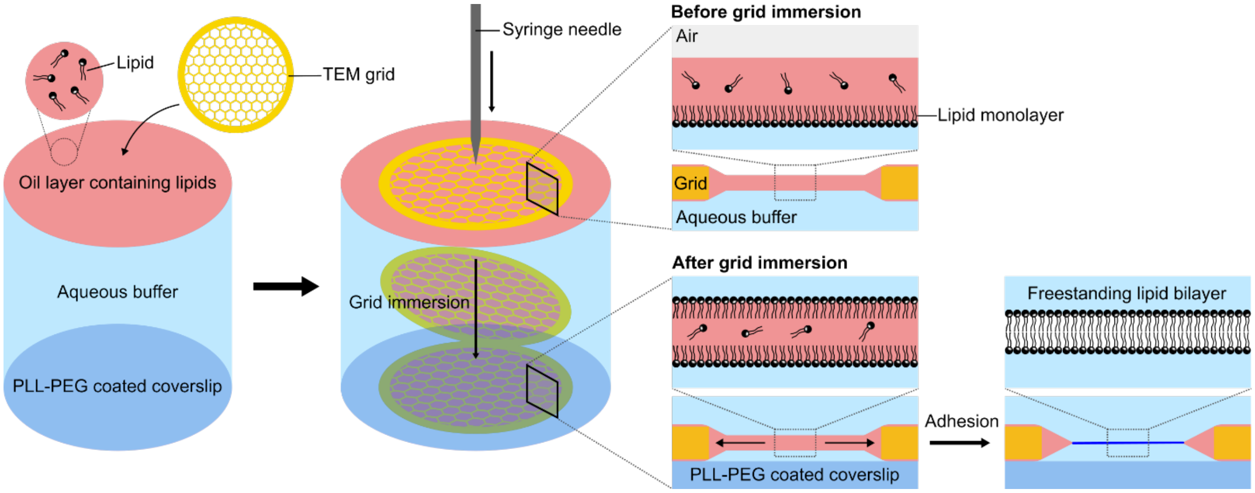


**Supplementary Figure 1. Schematic of freestanding planar lipid bilayer formation process.** Initially, a TEM grid with hexagonal holes was placed onto an oil layer containing lipids, leading to formation of a thin oil film within each grid hole with a lipid monolayer at the oil/buffer interface (before grid immersion). Next, the grid was immersed into the aqueous buffer using a syringe needle. Then, another lipid monolayer was formed at the oil/buffer interface. As the oil film in the grid hole became thinner, owing to oil draining back to the grid, spontaneous adhesion between the two lipid monolayers occurred, leading to a freestanding lipid bilayer (After grid immersion).

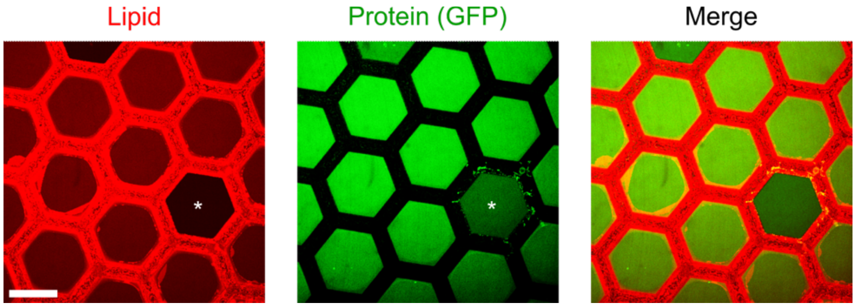


**Supplementary Figure 2. Homogeneous binding of GFP to suspended planar membranes.** Representative microscopic images after the addition of 1 μM of his-GFP to the membrane. Note that the fluorescence intensity of a vacant hole (marked with asterisk) is weaker than the surrounding membranes both in the lipid and protein channels. Membrane composition: 85 mol% DOPC, 15 mol% DGS-Ni-NTA, and 0.5 mol% Texas Red-DHPE. Buffer: 25 mM HEPES, 200 mM NaCl, pH 7.4. Scale bar, 100 μm.

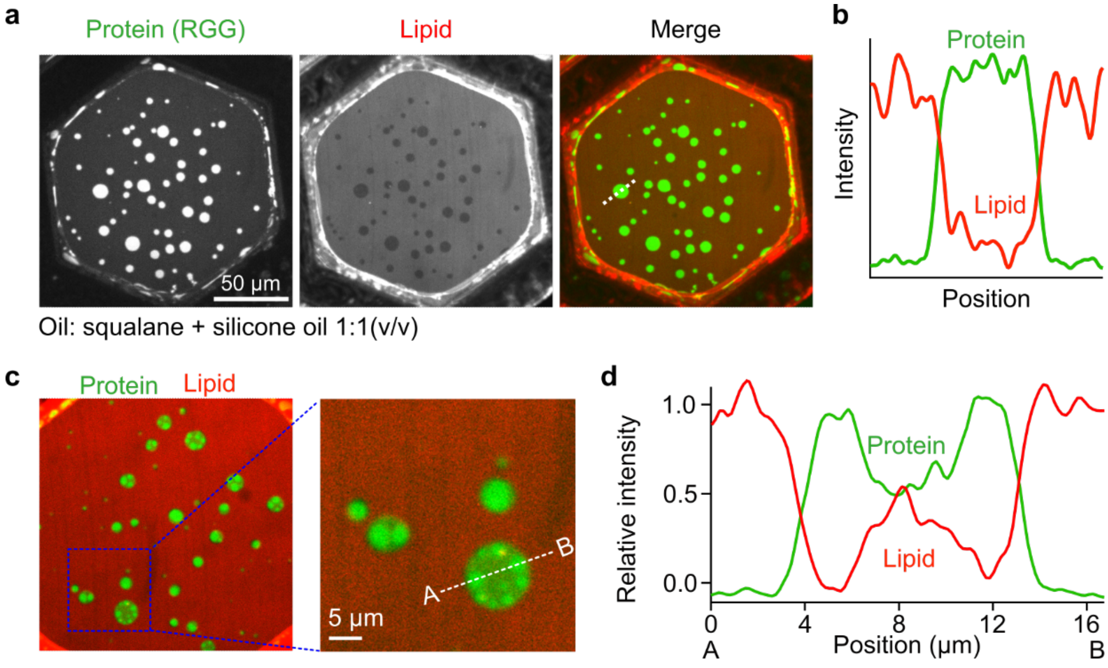


**Supplementary Figure 3. Lipid probe partitioning and transbilayer domain coupling of RGG condensates on solvent-free lipid membranes. a,** Representative images after the addition of 1 µM of his-RGG, labeled with Atto 488, to the membrane. The membrane could be considered as having an essentially solvent-free hydrophobic core, since squalane ($C_{30}H_{62}$), a hydrocarbon with a bulky structure, which does not easily enter the bilayers, was used instead of hexadecane ($C_{16}H_{34}$). Scale bar, 50 µm. **b,** Fluorescence intensity profile along the dotted white line in the merged channel in **a**, where green and red lines represent the intensity from the protein and lipid channels, respectively. **c,** Representative image showing transbilayer domain coupling. A region of interest within the blue dotted square is magnified. Scale bar, 5 µm. **d,** Intensity profiles along the dotted white lines in the magnified image in **c**. Membrane composition: 85 mol% DOPC, 15 mol% DGS-Ni-NTA, and 0.5 mol% Texas Red-DHPE. Buffer: 25 mM HEPES, 100 mM NaCl, pH 7.4.

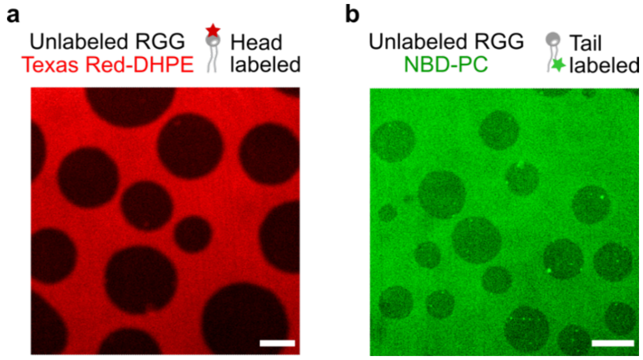


**Supplementary Figure 4. Lipid probe partitioning also occurs when condensates consist of unlabeled RGG.** Representative microscopic images in lipid probe channels, Texas Red-DHPE (**a**) and NBD-PC (**b**), after the addition of 1 µM of unlabeled his-RGG. Membrane composition: 85 mol% DOPC, 15 mol% DGS-Ni-NTA and 0.5 mol% lipid probe. Buffer: 25 mM HEPES, 50-100 mM NaCl, pH 7.4. Scale bars, 10 µm.

**Supplementary Movie 1.** Protein phase separation on several hexagonal lipid membranes over time. 1 µM of his-RGG labeled with Atto 488 was added. Time on the upper left corner represents the elapsed time since protein addition. Membrane composition: 75 mol% DOPC, 25 mol% DGS-Ni-NTA, 0.5 mol% Texas Red-DHPE. Buffer: 25 mM HEPES, 100 mM NaCl, pH 7.4. Scale bar: 50 µm.

**Supplementary Movie 2.** Fusion events between protein-rich domains on the membrane over time. 1 µM of his-RGG labeled with Atto 488 was added. Membrane composition: 85 mol% DOPC, 15 mol% DGS-Ni-NTA, 0.5 mol% Texas Red-DHPE. Buffer: 25 mM HEPES, 100 mM NaCl, pH 7.4. Scale bar: 10 µm.

**Supplementary Movie 3-5.** Dynamic domain fusions (Movie 3) and domain coupling **(Movie 4,5)** when protein-rich domains at different side of the membranes cross paths. 1 µM of his-RGG labeled with Atto 488 was added. Membrane composition: 75 mol% DOPC, 25 mol% DGS-Ni-NTA, 0.5 mol% Texas Red-DHPE. Buffer: 25 mM HEPES, 100 mM NaCl, pH 7.4.
